# Trophoblast stem cells and syncytiotrophoblasts lack inflammatory responses to LPS but retain robust interferon-mediated antiviral immunity

**DOI:** 10.1101/2025.10.23.684222

**Authors:** Cristine R. Camp, Joshua Baskaran, Matthew Brown, Carly Parker, Paige Drotos, Rachel C. West

**Affiliations:** Anatomy, Physiology, Pharmacology Department, College of Veterinary Medicine and Biomedical Science, Auburn University, Auburn AL, 36849

**Keywords:** trophoblast stem cell, syncytiotrophoblast, LPS, interferon beta, innate immune response, sex differences

## Abstract

Early pregnancy requires a tightly regulated pro-inflammatory environment shared between the primitive placenta and decidua. While immune balance supports successful implantation and placental invasion, disruptions in immune signaling during this period can impair implantation and lead to embryo loss. In this study, we investigated the molecular mechanisms underlying immune imbalance during implantation using a trophoblast stem cell (TSC) model. TSCs were cultured in either stem cell or syncytiotrophoblast (STB) differentiation medium and treated with either lipopolysaccharides (LPS) or interferon beta (IFNB). RT-qPCR and Western blotting revealed that LPS failed to induce a pro-inflammatory cytokine response in TSCs or STBs. In contrast, IFNB triggered a strong antiviral response in both TSCs and STBs. RNA-sequencing of IFNB-treated TSC and STB 3D spheroids revealed subtle differences between the TSCs and STB responses to interferons. Both TSC and STB IFNB-treated spheroids mount an interferon-mediated antiviral response; however, STB spheroid genes associated with the type I interferon response, viral RNA/DNA sensing, and antigen processing were upregulated. We also compared the interferon response between the CT27 (female) and CT29 (male) TSCs and STBs. While STBs showed minimal differences, the CT29 TSCs exhibited a markedly stronger interferon response than the CT27 TSCs. Collectively, these findings suggest that the primitive placenta is selectively responsive to interferon signaling rather than direct pathogen-associated stimuli. This implies that maternal immune activation, rather than microbial invasion, likely drives that placental immune response and embryo success at this stage. Understanding these dynamics underscores the importance of the maternal immune balance in early pregnancy success.

## Introduction

Early pregnancy requires the maintenance of a delicate balance of maternal immune signaling. At the onset of implantation, the fertilized egg has already reached the blastocyst stage and consists of the inner cell mass and the trophectoderm; the two cell lineages that will eventually give rise to the fetus and placenta, respectively. To initiate implantation, the trophectoderm adheres to then breaches the epithelial layer of the decidualized endometrium (Ochoa-Bernal & Fazleabas, 2020). Immediately following attachment to the decidualized endometrium, the trophectoderm fuses to form a highly invasive primitive syncytium that allows the embryo to breach the surface epithelium and invade into the underlying endometrium (Turco & Moffett, 2019). Both the pre-implantation embryo and invasive primitive placenta help stimulate an immunological cascade coming from the immune cells that comprise the decidua (Mor, Aldo, & Alvero, 2017). High levels of pro-inflammatory cytokines are necessary to support adequate trophoblast invasion and angiogenesis during early pregnancy (Sehring, Beltsos, & Jeelani, 2022). However, dysregulated inflammatory responses driven by infection or autoimmunity can lead to impaired implantation and embryo loss (Mor et al., 2017). The peri-implantation period is a notoriously difficult time to study due to the scientific and ethical limitations surrounding human embryos and early pregnancy. Therefore, there is a gap in our understanding as to how the cells of the primitive placenta interact with the uterine immune milieu in both normal and pathological circumstances.

Human trophoblast stem cells (TSCs) (Okae et al., 2018) have recently become a widely used model to study early placental development. TSCs derived from both the trophectoderm of blastocysts as well as villous column cytotrophoblast cells have similar transcriptome and DNA methylome signatures in all lines (Okae et al., 2018), suggesting the possibility of their use to study the formation of the primitive placenta. Further, the transcriptomes of TSCs compared to extended embryo culture, peri-implantation placental cells demonstrate similarities between TSCs and gestational day 8 cytotrophoblast cells (Logsdon et al., 2024), suggesting that TSCs model the early primitive cytotrophoblast. Additionally, syncytiotrophoblasts (STB) differentiated from TSCs are transcriptionally similar to the early STB described in gestational day 10 and 12 primitive placental cells (Logsdon et al., 2024). Another study documented that differentiating TSCs towards the syncytiotrophoblast (STB) led to the formation of an invasive STB capable of breaching the epithelium (Ruane et al., 2022), which is a hallmark of the primitive syncytium. Collectively, these data support the idea that TSCs are a novel model system capable of recapitulating early implantation dynamics.

Previous research using murine TSCs demonstrated that TSCs have attenuated immune responses to the inflammatory cytokines TNF_α_ and IFN_γ_ (Fendereski, Ming, Jiang, & Guo, 2024), suggesting that the primitive placenta can limit the cytotoxicity associated with a pro-inflammatory uterine environment. *Fendereski et al.* propose that the early embryo is an “immune-privileged structure” capable of growing and developing normally in a pro-inflammatory environment. While the peri-implantation stage embryo has an attenuated immune response to inflammatory cytokines, type I interferon and type II interferon receptors are present in the primitive placenta, with some receptors appearing as early as eight days post-fertilization (West et al., 2019). Thus, suggesting the embryo is capable of responding to interferon signaling from the endometrium and can mount an inflammatory response.

We aimed to characterize other inflammatory stimuli that would reflect a dysregulated uterine environment. As the peri-implantation stage placenta has type I interferon receptors (IFNAR), we hypothesized that the type I interferon, interferon beta (IFNB), could stimulate the trophoblast innate immune response. The role of IFNB during pregnancy is still unclear. One hypothesis asserts that low, constitutive levels of IFNB secreted by commensal bacteria within the reproductive tract are necessary for the maintenance of a healthy pregnancy (Ding et al., 2022). However, overproduction of IFNB due to a viral infection or autoimmune disorder can have detrimental effects on the growth and development of the placenta and fetus (Bundhun, Soogund, & Huang, 2017; Yockey et al., 2018).

We also aimed to assess the effects of lipopolysaccharides (LPS) on TSCs. LPS is a bacterial endotoxin that can induce a strong inflammatory response in other cell types, including the mature placenta. High levels of LPS can also have detrimental effects on placental development and fetal health (Brown, Maubert, Anton, Heiser, & Elovitz, 2019; Duval et al., 2019; Fan et al., 2019; Fricke et al., 2018); however, there is evidence to suggest that neither murine nor human embryonic stem cells (ESCs) mount an immune response to LPS (D’Angelo et al., 2017). We aimed to determine if placental cells representative of a peri-implantation embryo also have attenuated responses to LPS.

Two of the most commonly used TSC cell lines are genetically male and female. In this study we used the CT27 (female) and CT29 (male) TSC lines to uncover the molecular mechanisms behind the sexual dimorphism of early embryonic development and the placental innate immune response during pregnancy. There are inherent differences in fetal growth and metabolism between males and females (Campbell, Lucic Fisher, Brandon, Senior, & Bell-Anderson, 2022; Wells, 2007). However, we are now understanding that sex influences the placental innate and adaptive immune response during pregnancy (Baines & West, 2023). This sexual dimorphism leaves male fetuses vulnerable to miscarriage and stillbirth (Hassold, Quillen, & Yamane, 1983; Trudell, Cahill, Tuuli, Macones, & Odibo, 2015) as well as placental pathologies including gestational diabetes and late-onset preeclampsia (Broere-Brown et al., 2020; Cooperstock & Campbell, 1996; Huppertz, 2008). Interestingly, even with adverse pregnancy outcomes skewing male, the primary sex ratio at birth is slightly male-biased (Orzack et al., 2015). We used a 3-dimensional (3D) spheroid model to assess the differences in immune stimulated CT27 (female) and CT29 (male) TSC and STBs. Our results demonstrate inherent differences in trophoblast responses to inflammatory stimuli as well as differences in gene expression and innate immune response between cell lines.

## Materials and Methods

### Trophoblast Stem Cell Culture

TSCs were first generated by Okae et al. (Okae et al., 2018) and obtained through a material transfer agreement with the RIKEN Cell Engineering Division. Tissue culture treated plates were coated with Laminin-511 (iMatrix-511, ReproCell) and incubated at 37°C for 1 hour. TSCs were plated and cultured in TSC medium containing DMEM/F12 (Thermo Fisher, #11320082), 0.3% bovine serum albumin (BSA) (Fisher Scientific, #BP9704100), 1% ITS-X (Gibco, #51500-056), 0.2% fetal bovine serum (FBS) (Thermo Fisher, #16141002) 0.1 mM 2-mercaptoethanol, 50 ng/mL epidermal growth factor (EGF) (Millipore Sigma, #E9644), 1.5 µg/mL L-ascorbic acid (Millipore Sigma, #A8960), 2 µM CHIR99021 (ReproCell, #04-0004), 0.5 µM A83-01 (ReproCell, #04-0014), 1 µM SB431542 (ReproCell, #04-0010), 0.8 mM valproic acid (VPA) (Millipore Sigma, #P4543), and 5 µM Y27632 (ReproCell, #04-0012). TSCs were cultured at 37°C in 5% CO_2_. Once TSCs, reached approximately 70% confluency, TSCs were dissociated in TrypLE (Thermo Fisher, #12563011) for 12-14 minutes and passaged to a new iMatrix coated plate. To differentiate towards the STB, tissue culture treated plates were coated with 2.5 µg/mL of mouse Collagen IV (Corning, #354233). TSCs were plated on new wells and given 24 hours to recover in TSC medium. After 24 hours, TSCs were cultured in STB media conditions: DMEM/F12, 0.3% BSA, 1% ITS-X, 0.1 mM 2-mercaptoethanol, 205 µM Y27632, 2 µM Forskolin (Millipore Sigma, #F6886), and 4% knockout serum replacement (KSR) (Thermo Fisher, #10828010). Cells were cultured in STB medium for 5 days and medium was replenished every other day. To grow cells as 3D spheroids, tissue culture treated plates were rinsed with Anti-Adherence Rinse solution (Stem Cell Technologies, #07010). TSCs were grown in 3D stem conditions for 48 hours before IFNB treatment. STBs were grown in 3D STB conditions for 4 days before IFNB treatment. All cells were used between serial passages 23-28 and care was made that cells at the same passage number were used within experiments. The absence of mycoplasma was confirmed monthly using the MycoStrip mycoplasma detection kit (Invivogen, #rep-mys-50).

### Fibroblast Cell Culture

Human primary dermal fibroblasts (HDFs) were purchased from ATCC (PCS-201-012). HDFs were cultured in fibroblast basal medium (ATCC) supplemented with a fibroblast growth kit – low serum (ATCC, #PCS-201-041) containing 7.5 mM L-glutamine, 5 ng/mL rh fibroblast growth factor (FGF), 5 µg/mL rh insulin, 1 µg/mL hydrocortisone, 50 µg/mL ascorbic acid, and 2% FBS. Cells were cultured to 70% and passaged as needed.

### LPS and IFNB Treatment

TSCs, STBs, and HDFs were treated with 250 ng/mL LPS (Sigma, #L4391) for 6 hours for RT-qPCR experiments or for 15, 30, and 60 minutes and 24 hours for Western blotting. Cells were treated with 250 U/mL IFNB (R&D Systems, #8499) for the same time points.

### RNA Isolation and RT-qPCR

All RT-qPCR experiments were designed and run in concordance with the MIQE guidelines (Bustin et al., 2009). RNA was isolated from cells using a Qiagen RNeasy mini kit (#74104) following manufacturer’s instructions. Complementary DNA (cDNA) was generated from 500 ng total RNA using the qScript cDNA synthesis kit (QuantaBio, 95047). After generation of cDNA, cDNA was diluted to 50 ng per reaction and mixed with PerfeCTa SYBR Green FastMix (QuantaBio, 101414). Primer sequences for primers used in RT-qPCR reactions are displayed in Table 1. For RT-qPCR, reactions were incubated at 95°C for 10 minutes then underwent 40 cycles of 95°C for 15 seconds and 60°C for 1 minute using the QuantStudio 5 PCR system (ThermoFisher). Each reaction was conducted in duplicate. The averages of the technical duplicates were used to normalize relative expression against GAPDH. Fold change was determined against an untreated control for each cell type.

**Table 1.**
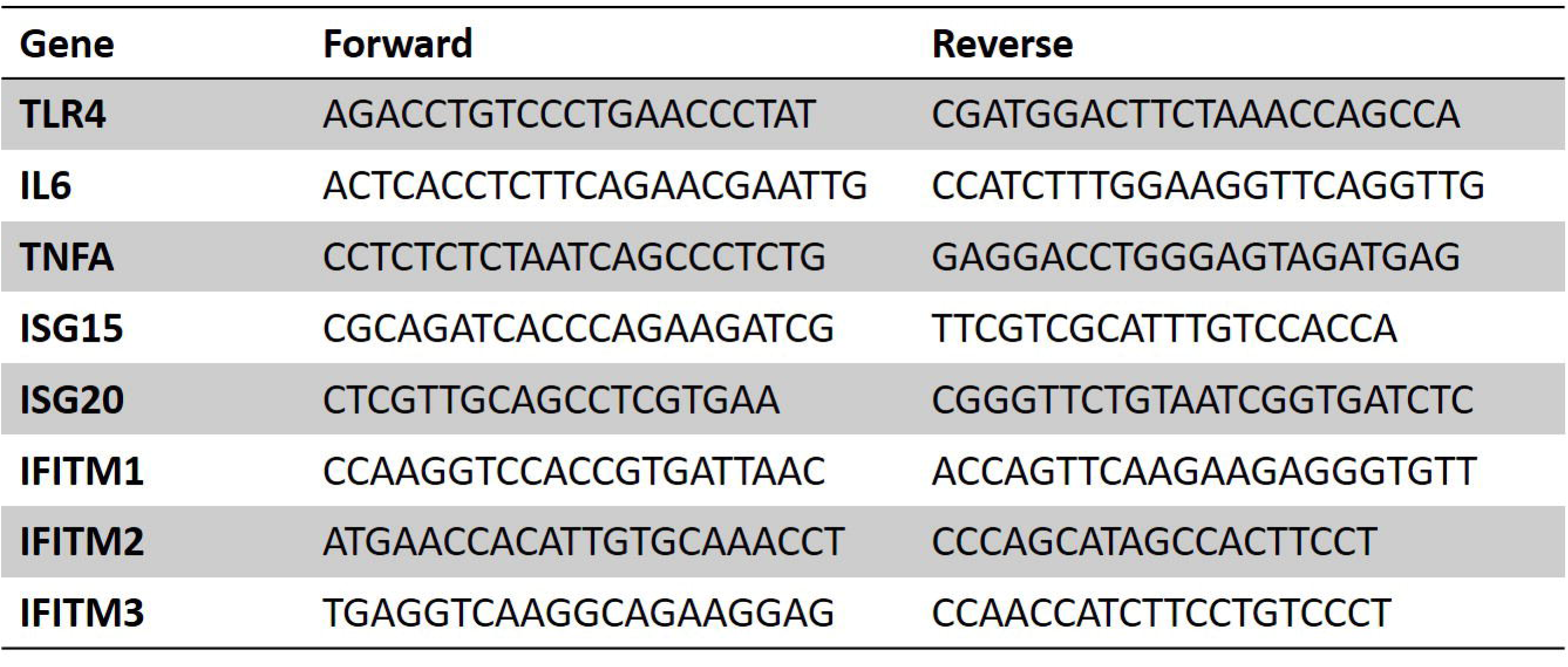
Primers used for RT-qPCR.

### Western blotting

Cells were lysed in RIPA buffer (Pierce, ThermoFisher, #89900) with Halt phosphatase inhibitor cocktail (ThermoFisher, #78442), and cOmplete mini EDTA Free protease inhibitor (Millipore Sigma, #11836170001). After lysis, protein was sonicated in 3 cycles of 15s on/30s off. Total protein was quantified using a Qubit Broad Range Protein Assay Kit (ThermoFisher, #A50668). After quantification, protein was diluted to 20 µg using 4X Laemmli buffer (BioRad, #1610747) and 40X dithiothreitol (DTT) (Millipore Sigma, #10708984001). Protein was denatured for 10 minutes then allowed to cool to room temperature for 5 minutes. After denaturation, protein was electrophoresed in a 4-20% mini-protean TGX stain-free protein gel (Bio-Rad, #4568096). Protein was then transferred for 5 minutes using the Trans-Blot Turbo transfer system onto a nitrocellulose membrane (BioRad, #1704156). After protein transfer, membranes were blocked in 3% BSA in 1X TBS-T (BioRad, #1706435) then incubated with primary antibodies (pERK, Cell Signaling Technology, #4370; STAT1, Cell Signaling Technology, #9172; TLR4, Santa Cruz Biotechnology, sc293072) overnight at 4°C. After incubation, membranes were washed three times in TBS-T then incubated for 1 hour at room temperature with an infrared dye-conjugated secondary antibody. Membranes were imaged using a LICORbio Odyssey imaging system. Beta-Actin (ACTB) (Cell Signaling Technology, #3700S) or GAPDH (Cell Signaling Technology, #2118S) were used to normalize protein in cell lysates. Densitometry semi-quantitative analysis was performed using ImageJ imaging software and relative density was calculated as a percentage of control protein after normalization.

### RNA-sequencing

Purified RNA was sent to Novogene for library construction and sequencing. Briefly, messenger RNA (mRNA) was extracted from total RNA by poly-T oligo-attached magnetic beads. mRNA was subject to library preparation and cDNA was multiplexed by PCR amplification. Libraries were quantified using Qubit and size distribution analyzed using the Bioanalyzer 2100 system (Agilent). Quantified libraries were pooled and sequenced using the Illumina NovaSeq X Plus platform.

Raw fastq reads were trimmed to remove adapters and low-quality reads using fastp. Clean reads were mapped to the human genome (hg38) and individual mapped reads were adjusted to provide fragments per kilobase per transcript sequence per millions base pairs sequenced (FPKM). Differential expression gene analysis was performed using the DESeq2 R package (1.20.0). Genes were deemed differentially expressed if they provided a false discovery rate of <0.05 and a log2foldchange > 1 and < -1. Comparison of data sets (i.e. TSCs vs. IFN-TSCs and STBs vs. IFN-STBs) to determine differential gene expression between data sets was performed using the DESeq2 R package (1.20.0). Genes were determined to be preferentially enriched in a cell type if they had an adjusted p-value (padj) <0.05 and a log2foldchange >1 or <- 1. Gene ontology (GO) enrichment analysis of differentially expressed genes was performed using the clusterProfiler R package. GO terms with padj < 0.05 were considered significantly enriched.

### Statistics

For molecular experiments, at least three, independent biological replicates were used for each experiment and experiments were repeated at least three times. Statistical significance was evaluated with ANOVA with Tukey’s test for multiple comparisons. All statistics were generated using GraphPad Prism 9 software. P-values less than 0.05 were considered statistically significant.

## Results

### LPS induces a limited inflammatory response in TSCs and STBs compared to IFNB

We first measured the mRNA levels of *TLR4* and the pro-inflammatory cytokines interleukin-6 (*IL6*) and tumor necrosis factor alpha (*TNFA*). We also measured mRNA levels of the interferon stimulated genes (ISGs), *ISG15* and *ISG20* as well as the interferon inducible transmembrane genes (IFITM), *IFITM1, IFITM2,* and *IFITM3.* While LPS triggered a significant increase in *IL6* and *TNFA* mRNA levels in the HDF, LPS did not stimulate an increase in *TLR4, IL6,* or *TNFA* in TSCs (Fig 1A-C). Alternatively, TSC treatment with IFNB induced a robust response in the ISGs, *ISG20, IFITM1, IFITM2,* and *IFITM3* (Fig 1E-H) and an elevated, albeit not significantly, response in *ISG15* levels (Fig 1D) compared to wild type and LPS treated TSCs.

**Figure 1.**
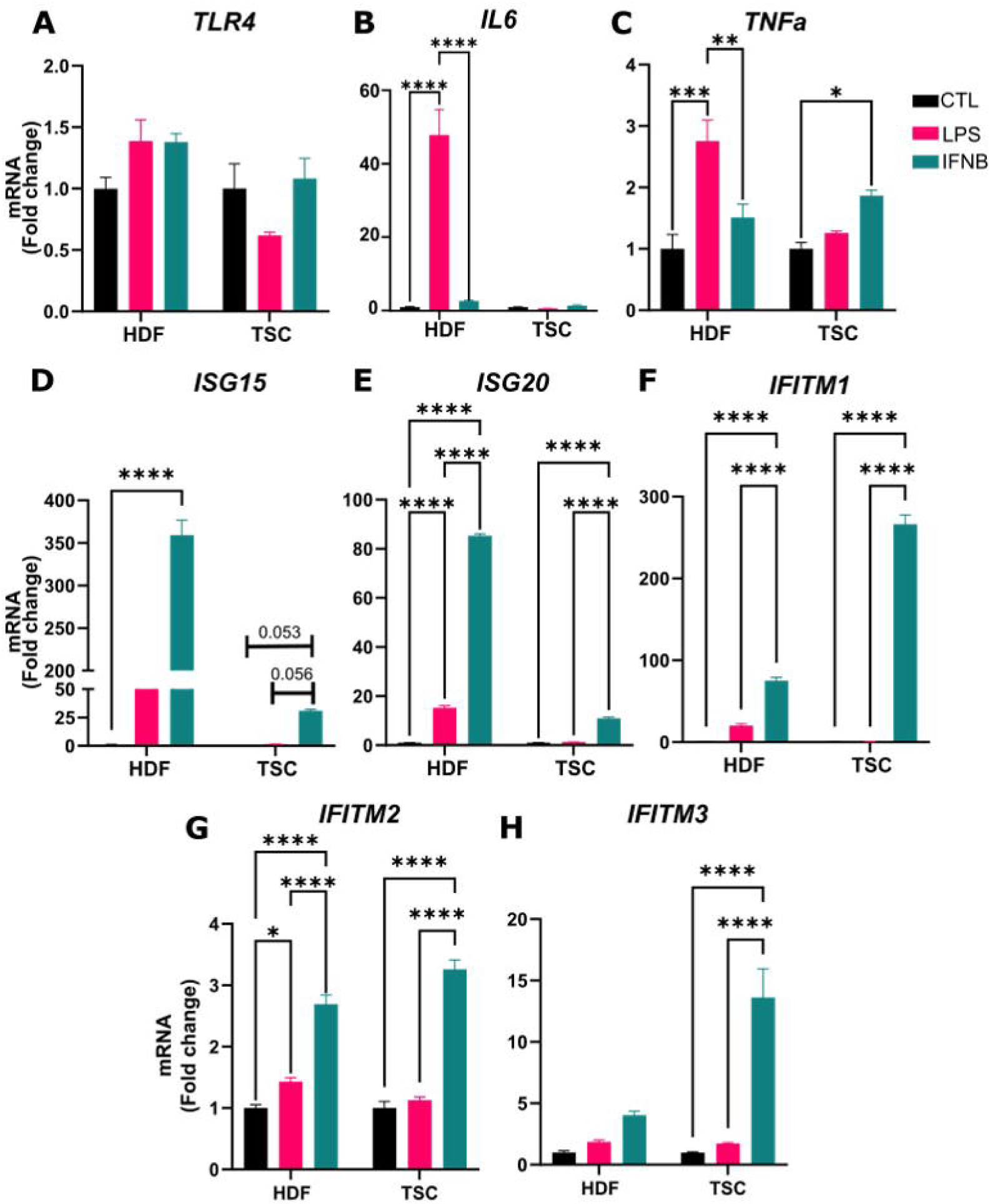
Pro-inflammatory cytokine and ISG mRNA levels in LPS and IFNB-treated TSCs. While there is no increase in **A)** pattern recognition receptor *TLR4,* LPS treatment significantly increased mRNA levels of the **B)** pro-inflammatory cytokine *IL6* and **C)** pro-inflammatory cytokine *TNFa* in HDFs but not TSCs. However, LPS treatment did stimulate a modest, but significant increase in *TNFa* in the TSCs. Alternatively, IFNB treatment induced an ISG response in the HDFs and TSCs for **D)** *ISG15* **E)** *ISG20* **F)** *IFITM1* **G)** *IFIMT2.* Only the TSCs had increased levels of **H)** *IFITM3* in response to IFNB. * indicates p < 0.05, ** indicates p < 0.01, *** indicates p < 0.005, and **** indicates p < 0.001.

LPS also failed to stimulate a pro-inflammatory or IFN response in STBs (Figure 2A-H). However, IFNB induced a robust response in both the STBs and HDF. *TLR4* mRNA levels were modestly but significantly higher in IFNB treated STBs compared to LPS treated (Figure 2A). Additionally, *TNFA* was also modestly but significantly higher in IFNB treated TSCs and STBs compared to control and LPS treated STBs. *ISG20, IFITM1, IFITM2,* and *IFITM3* were all significantly elevated in IFNB treated STBs. These results suggest that LPS does not stimulate a pro-inflammatory response in TSCs or STBs. However, IFNB can stimulate a robust IFN response in both cell types.

**Figure 2.**
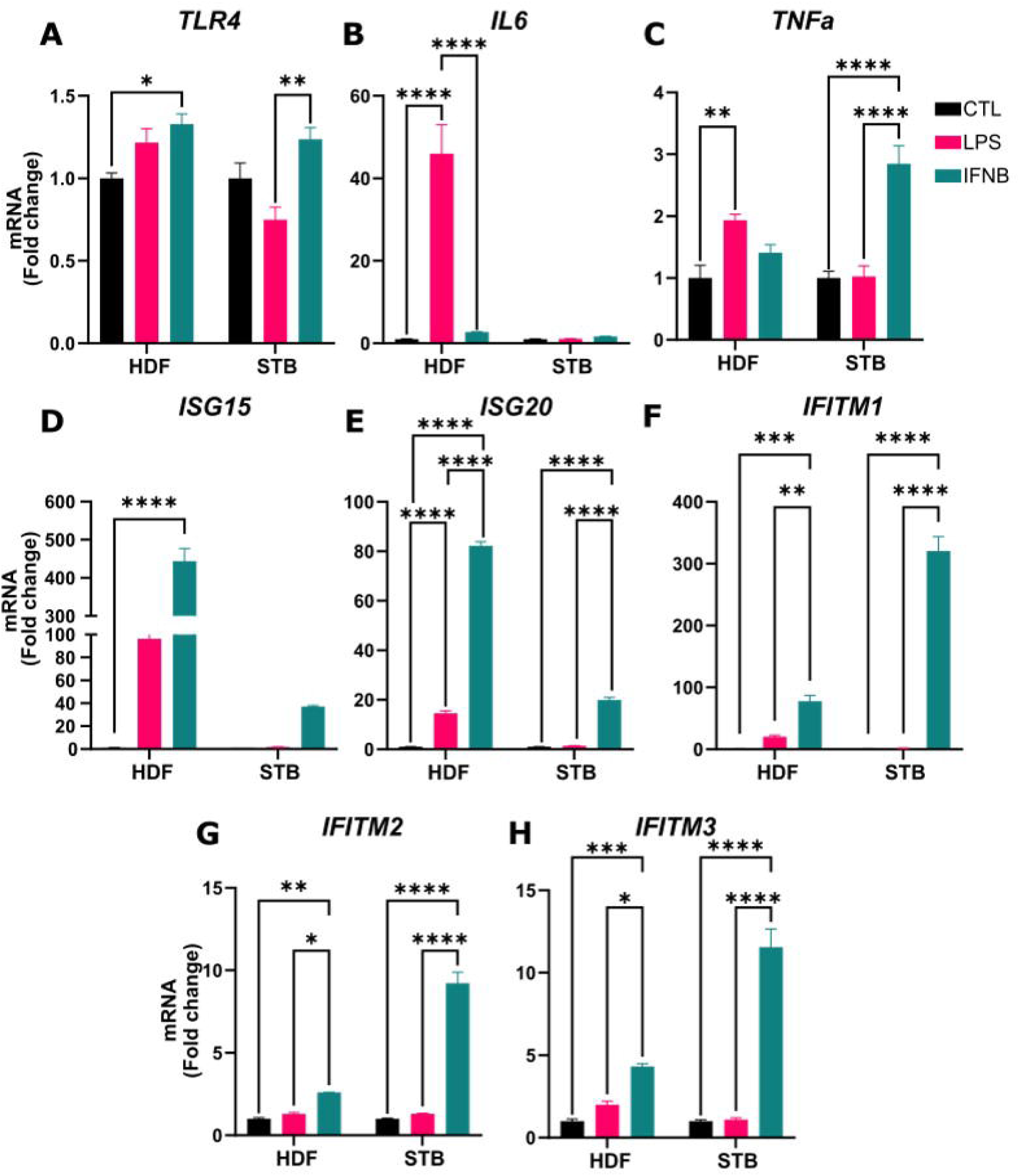
Pro-inflammatory cytokine and ISG mRNA levels in LPS and IFNB-treated STBs. LPS treatment did not induce a significant change in the **A)** pattern recognition receptor *TLR4.* However, IFNB treatment did trigger a modest but significant increase. LPS treatment significantly increased mRNA levels of the **B)** pro-inflammatory cytokine *IL6* and **C)** pro-inflammatory cytokine *TNFa* in HDFs but not STBs. *TNFa* levels were significantly elevated in STBs treated with IFNB. IFNB treatment also induced an ISG response in the HDFs for **D)** *ISG15.* The ISGs, **E)** *ISG20* **F)** *IFITM1* **G)** *IFIMT2* and **H)** *IFITM3* were significantly increased in both the HDFs and STBs. * indicates p < 0.05, ** indicates p < 0.01, *** indicates p < 0.005, and **** indicates p < 0.001.

We next used Western blotting to determine protein levels of TLR4 in the unstimulated HDF, TSCs, and STBs and found no differences in protein abundance between the three cell types (Figure 3A). To assess if LPS stimulates phosphorylation of ERK1/2 in TSCs and STBs, we treated cells with LPS then collected protein 15, 30, and 60 minutes after treatment. There was a delayed, modest increase of pERK in both the TSCs and STBs. Conversely, there was a robust increase to LPS at 15 minutes followed by a rapid decrease of pERK in the HDF control (Figure 3B). We also used Western blotting to demonstrate that IFNB treatment increased total STAT1 protein in HDF, TSCs, and STBs after 24 hours of treatment (Figure 3C). These data further demonstrate that Type I interferons trigger a robust antiviral response in TSCs and STBs, whereas LPS does not stimulate the production of pro-inflammatory cytokines in TSCs or STBs.

**Figure 3.**
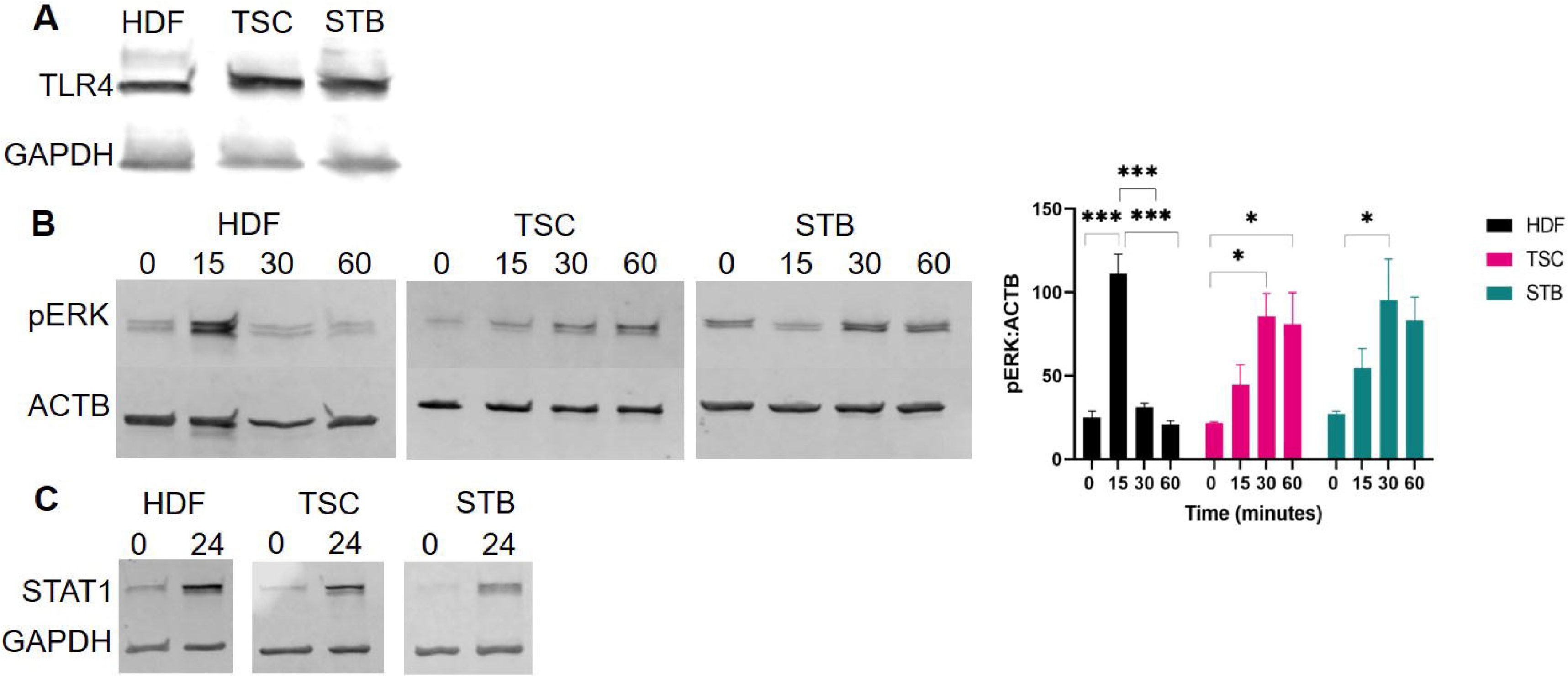
LPS triggers rapid phosphorylation of ERK in HDFs but not TSCs or STBs. Total STAT1 is increased in all cell types after IFNB treatment for 24 hours. **A)** There are no differences in protein levels for TLR4 between the untreated HDF, TSCs, and STBs. **B)** Protein levels for pERK 0, 15, 30, and 60 minutes after LPS treatment in HDFs, TSCs, and STBs. There was a significant increase in pERK 15 minutes post-LPS treatment followed by a significant decrease 30 and 60 minutes post-treatment. Alternatively, there was a modest but significant increase in pERK 30 and 60 minutes post-LPS treatment in TSCs and no significant increase in pERK at any time point compared in the STBs compared to the untreated control. **C)** Proteins levels for total STAT1 after 24 hours of IFNB treatment. * indicates p < 0.05, *** indicates p < 0.005.

### IFNB elicits a robust antiviral response in both TSCs and STBs

We next compared the transcriptomes of untreated and IFN-treated TSCs and untreated and IFN-treated STBs. Using RNA-seq, we analyzed untreated TSCs and STBs and IFN-treated TSCs and STBs. The cell types were first analyzed for gene expression and genes with padj values <0.05 and a log2foldchange >1 were designated as significant differentially expressed genes. There were 252 (245 upregulated, 7 downregulated) significantly different transcripts between the untreated TSCs and IFN-treated TSCs (Figure 4A, Supplemental File 1). Similarly, there were 185 DEGs (163 upregulated, 22 downregulated) between the untreated STBs and IFN-treated STBs (Figure 4A, Supplemental File 2). GO analysis of IFNB-treated TSCs and STBs compared to their respective controls were very similar (Supplemental Figure 1), with GO terms related to response to virus and the type I interferon signaling pathway strongly associated with both IFNB-treated cell types. We next used GO analysis to compare differences between the TSCs and STBs and IFN-treated TSCs and IFN-treated STBs. Many pathways related to syncytialization, steroidogenesis, and protein processing were significantly enriched in the STBs and IFN-treated STBs compared to the TSCs and IFN-treated TSCs (Supplemental File 3). However, several top enriched terms for both the STBs and IFN-treated STBs compared to the TSCs and IFN-treated TSCs were related to the inflammatory response (Supplemental File 3); therefore, we next compared the DEG profiles between the TSCs and IFN-treated TSCs and STBs and IFN-treated STBs data sets to identify genes and pathways that were preferentially enriched only in the IFN-treated STBs and IFN-treated TSCs compared to the other cell types. This comparison revealed 26 genes preferentially enriched in the IFN-treated TSCs and 180 genes preferentially enriched in the IFN-treated STBs (Figure 4B, Supplemental File 4). Additionally, there were similar gene expression patterns for 5 genes were observed in both the IFN-treated TSCs and STBs (Figure 4B, Supplemental File 4). Clustering analysis was performed to visualize differences in gene expression between each treatment group (Figure 4C, Supplemental File 4).

**Figure 4.**
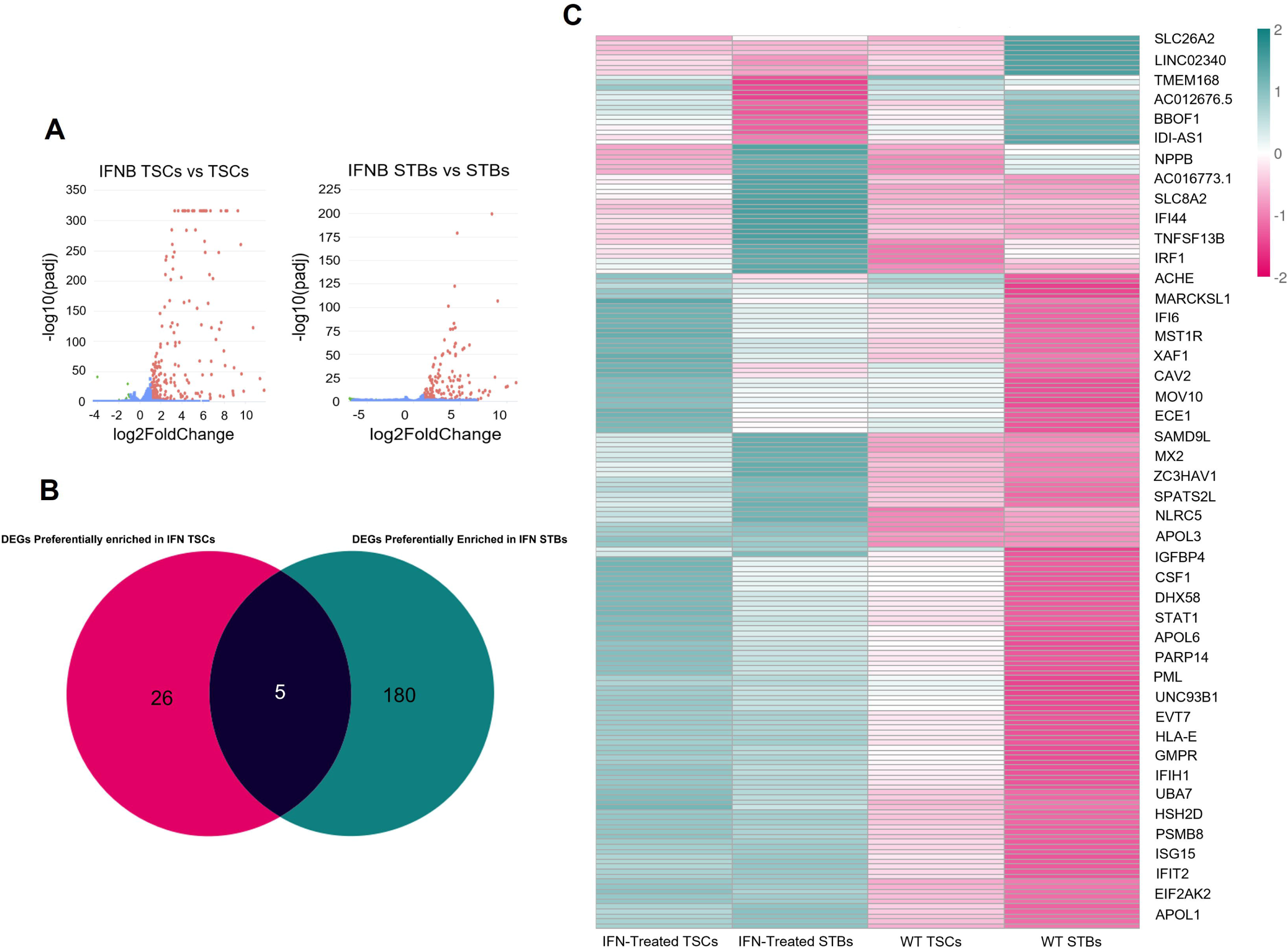
IFNB triggers an antiviral response in both TSCs and STBs but STBs also have increased transcript levels for pro-inflammatory cytokines and chemokines and genes for recruitment of immune cells. **A)** IFNB treated TSCs had 252 (245 upregulated, 7 downregulated) DEGs compared to untreated TSCs. IFNB-treated STBs had 185 (163 upregulated, 7 downregulated) DEGs compared to untreated STBs. Red points represent transcripts with a log2foldchange >1 and an adjusted p-value of < 0.05. Green points represent transcripts with a log2foldchange -1< and an adjusted p-value of <0.05. **B)** Venn diagram of DEGs preferentially enriched in the IFN-treated TSCs or IFN-treated STBs after comparison between the two data sets comparing treated cells against their respective controls. **C)** Heat map clustering of the preferentially enriched genes found in the IFN-treated TSCs and IFN-treated STBs. The color spectrum, ranging from teal to pink, indicates high to low normalized levels of expression of each gene.

GO analysis of the IFN-treated STB preferentially enriched DEGs revealed upregulation of GO tems related to response to type I interferon and response to virus (Table 2, Supplemental File 4). Many of the 180 preferentially enriched genes in the IFN-treated STBs were canonical ISGs (*STAT1, IRF1, IRF7, SOCS1, IFIT1, IFIT2, IFIT3, IFIT5, OAS1, OAS2, OAS3, ISG15, ISG20, IFI44, IFI6, IFI27, IFI35).* Additionally, genes related to viral RNA/DNA sensing and restriction machinery (*IFIH1, DDX58, TLR3, SAMHD1, ADAR, TRIM14, TRIM22, TRIM25, TRIM38, TRIM69)* and antigen processing and the MHC class I pathway (*HLA-C, HLA-E, HLA-F, TAP1, TAP2, NLRC5)* were upregulated in the IFN-treated STBs. Several other genes related to the inflammasome and immune cell activation (*CASP1, CASP8 TNFSF10, MYD88, CARD11)*, and cytokines and chemokines (*CXCL16, CCL28, TNSF13B, CSF1, PDCD1LG2)* were also elevated. This gene expression profile suggests a robust, coordinated interferon-driven antiviral and immune-activation program within the STBs. Alternatively, GO analysis revealed no upregulated GO terms within the preferentially enriched IFN-treated TSC DEGs. While there were some innate immune related DEGs (*C3AR1, PIK3AP1, IKZF1, SIGLEC6, LY63, TRIM56, DUSP10, FLG1)*, the gene expression profile was more representative of a heterogeneous pool of genes, suggesting a modest interferon response compared to the STBs.

**Table 2.**
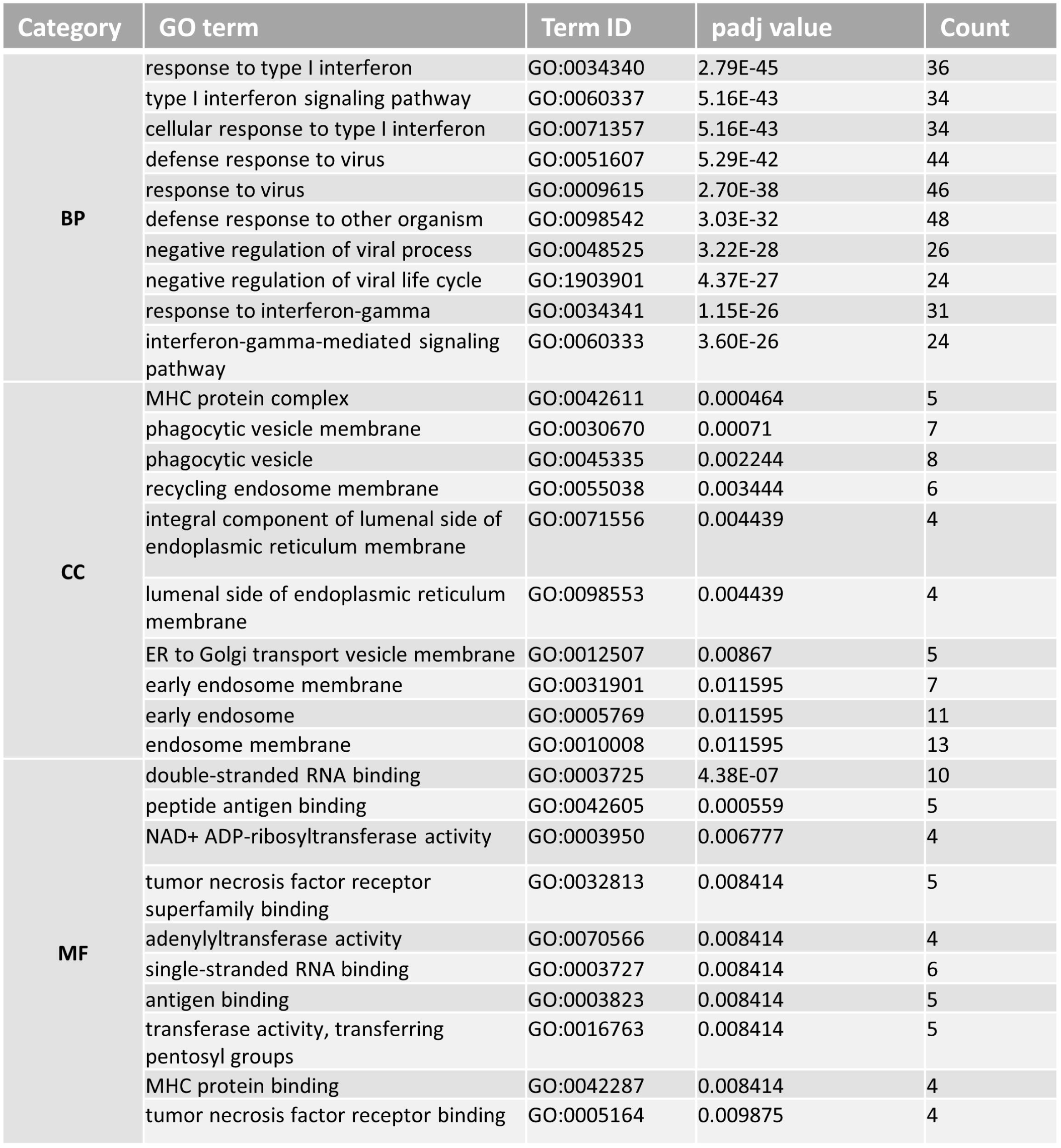
The top 10 significant GO terms for each of BPs, MFs, and CCs identified through analysis of the preferentially enriched IFN-treated STB DEGs. A comprehensive list of all GO terms can be found in Supplemental File 4.

### There are inherent genetic differences between TSC lines

The experiments described in Figures 1-4 were conducted using the CT29 TSC cell line; however, *Okae et al.* generated several TSC lines (Okae et al., 2018). As the CT29 TSCs are genetically XY, we aimed to compare the differences between XY TSCs against the XX CT27 cell line. Compared to the CT27 TSCs, the CT29 TSCs had 2,342 DEGs (1,221 upregulated, and 1,121 downregulated) (Figure 5A, Supplemental File 5). Additionally, there were 1,361 DEGs (647 upregulated, 714 downregulated) in the CT29 STBs compared to the CT27s (Figure 5A, Supplemental File 6). We next compared the gene profiles between the CT29 vs. CT27 TSC and CT29 and CT27 STB differential gene expression analyses to identify genes that were preferentially enriched in each cell subtype. This comparison revealed 1,791 genes preferentially enriched in the CT29 TSCs and the 810 genes preferentially enriched in the CT29 STBs. There were 551 shared genes between the CT29 TSCs and STBs compared to the CT27s (Figure 5B, Supplemental File 7).

**Figure 5.**
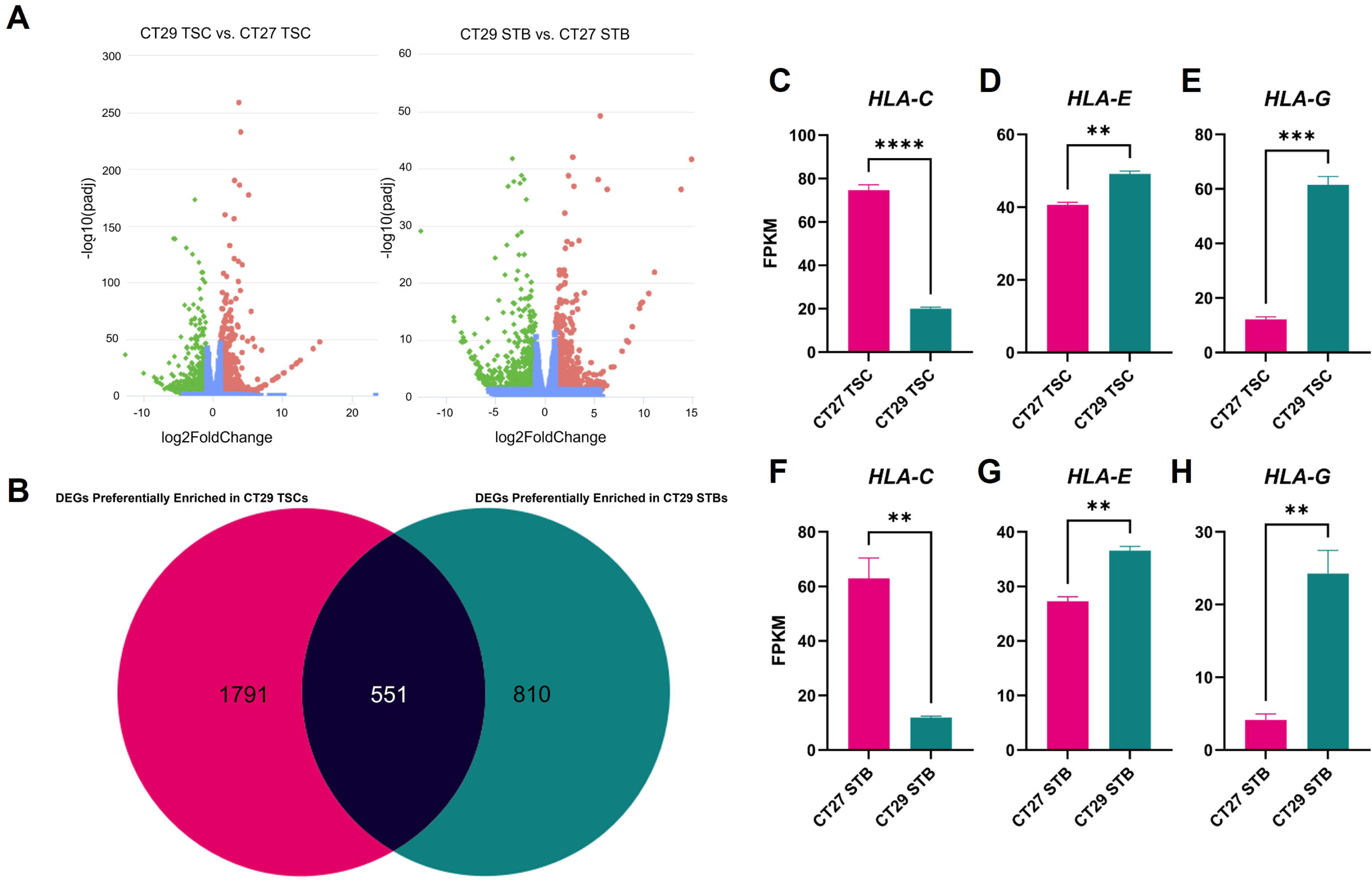
Wild type CT27 TSCs and STBs have differences in gene expression compared to CT29 TSCs and STBs. **A)** Volcano plots of DEGs. The CT29 TSCs had 1,221 upregulated and 1,121 downregulated DEGs compared to the CT27 TSCs. The CT29 STBs had 647 upregulated and 714 downregulated DEGs compared to the CT29 STBs. Red points represent transcripts with a log2foldchange >1 and an adjusted p-value of < 0.05. Green points represent transcripts with a log2foldchange -1< and an adjusted p-value of <0.05. **B)** Venn diagram of DEGs preferentially enriched in the CT29 TSCs or CT29 STBs after comparison between the two data sets comparing CT27 and CT29 TSCs and CT27 and CT29 STBs. **C-E)** FPKM values of *HLA-C, HLA-E,* and *HLA-G* between the CT27 and CT29 TSCs. **F-H)** FPKM values of *HLA-C, HLA-E* and *HLA-G* between the CT27 and CT29 STBs. Error bars represent SEM (n=3) calculated from RNA-seq analysis. ** indicates p<0.01, *** p<0.005, **** p<0.001

We performed GO analysis on the DEGs preferentially enriched in the CT29 and CT27 TSCs and STBs. There were several significant GO terms in the CT29 TSCs (Table 3, Supplemental File 7). Many of these terms were related to cell migration and extracellular matrix organization. Alternatively, when we performed GO analysis on the DEGs preferentially enriched in the CT27 TSCs, terms related to detoxification of metals were significantly upregulated (Table 4, Supplemental File 7). Many of the upregulated genes in the CT27 TSCs belong to the metallothionein (MT) family, including *MT2A, MT1G, MT1E, MT1H, MT1F,* and *MT1X* (Supplemental Figure 2) which are important for detoxification of metals and protection against oxidative stress (Thirumoorthy, Manisenthil Kumar, Shyam Sundar, Panayappan, & Chatterjee, 2007). There were no differences in gene expression of MTs between the CT27 and CT29 STBs.

**Table 3.**
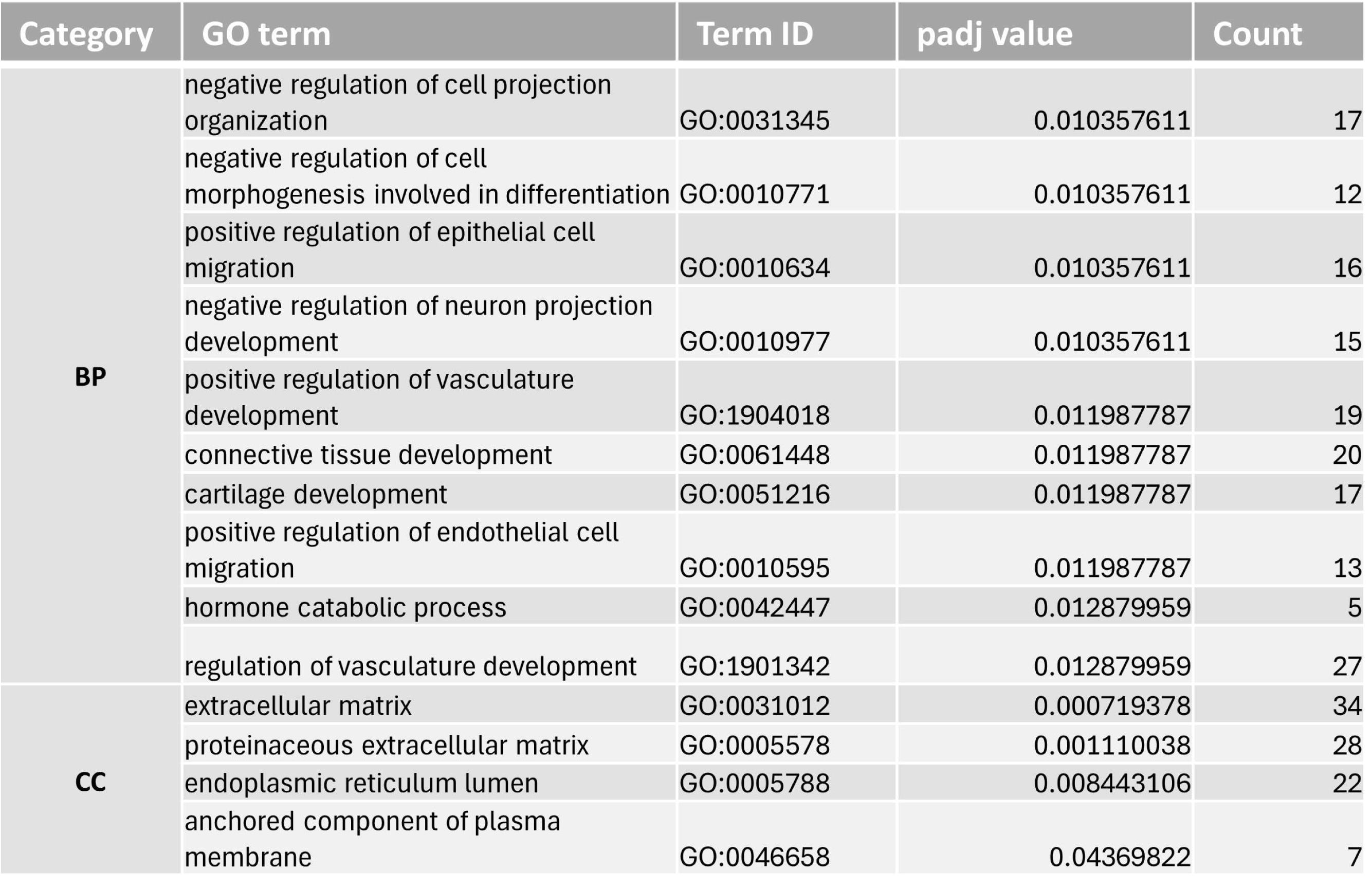
The top 10 significant GO terms for BP and all significant CC terms identified through analysis of the preferentially enriched CT29 TSC DEGs. A comprehensive list of the remaining significant terms can be found in Supplemental File 7.

**Table 4.**
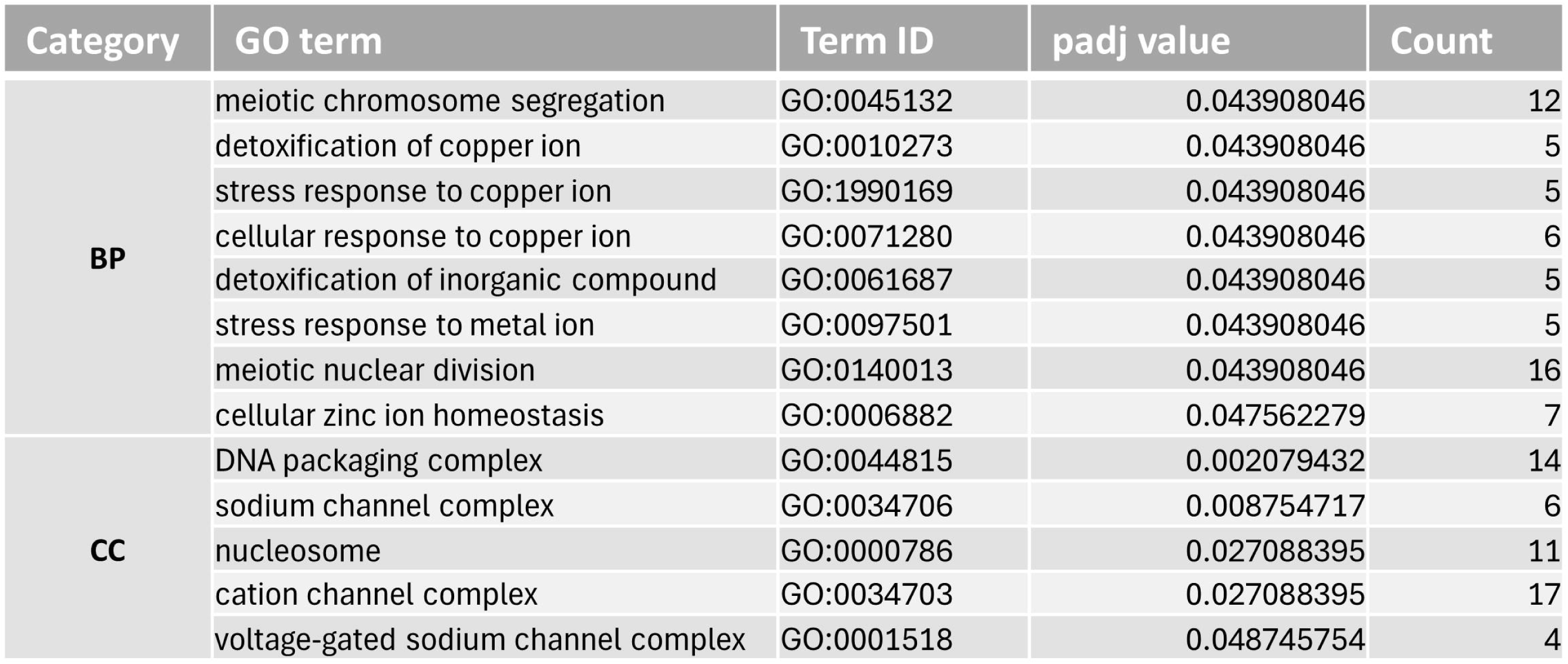
All significant GO terms identified through analysis of preferentially enriched CT27 TSC DEGs.

There were 11 significant pathways in the DEGs preferentially enriched in the CT29 STBs (Table 5, Supplemental File 7). Most of the terms enriched in the CT29 STBs were related to protein folding and processing. Many of the upregulated genes related to protein folding and processing belonged to the heat shock protein family (*HSPA1A, HSPA4L, HSPA1B, HSPE1, HSPH1)*. Alternatively, GO analysis of the DEGs preferentially enriched in the CT27 STBs revealed several pathways related to cholesterol metabolism and steroid biosynthesis (Table 6, Supplemental File 7). Protein folding, processing, and cholesterol metabolism and steroid biosynthesis are all hallmark pathways associated with the STB; however, it appears that the CT29s and CT27s are divergent, at least at the transcriptional level, in which pathways are prioritized upon differentiation.

**Table 5.**
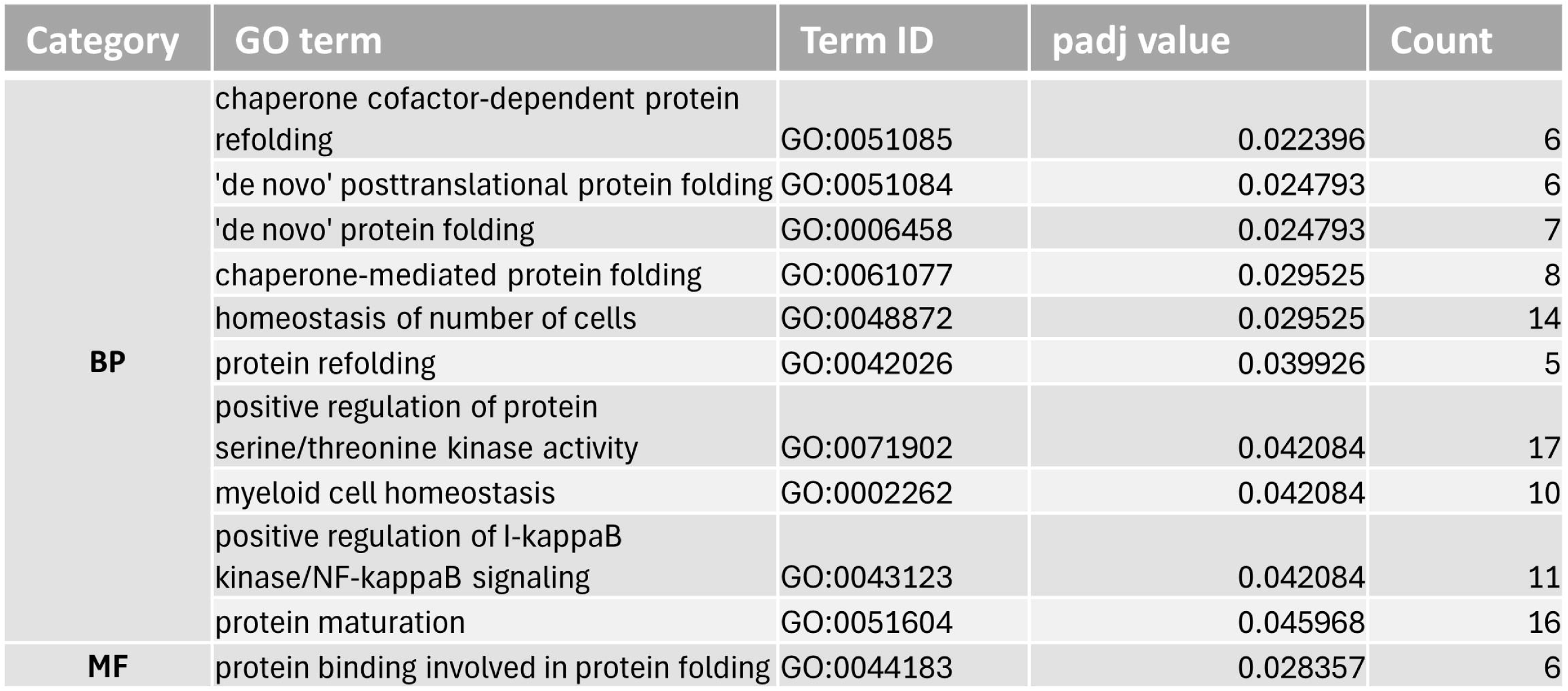
All significant GO terms identified through analysis of preferentially enriched CT29 STB DEGs.

**Table 6.**
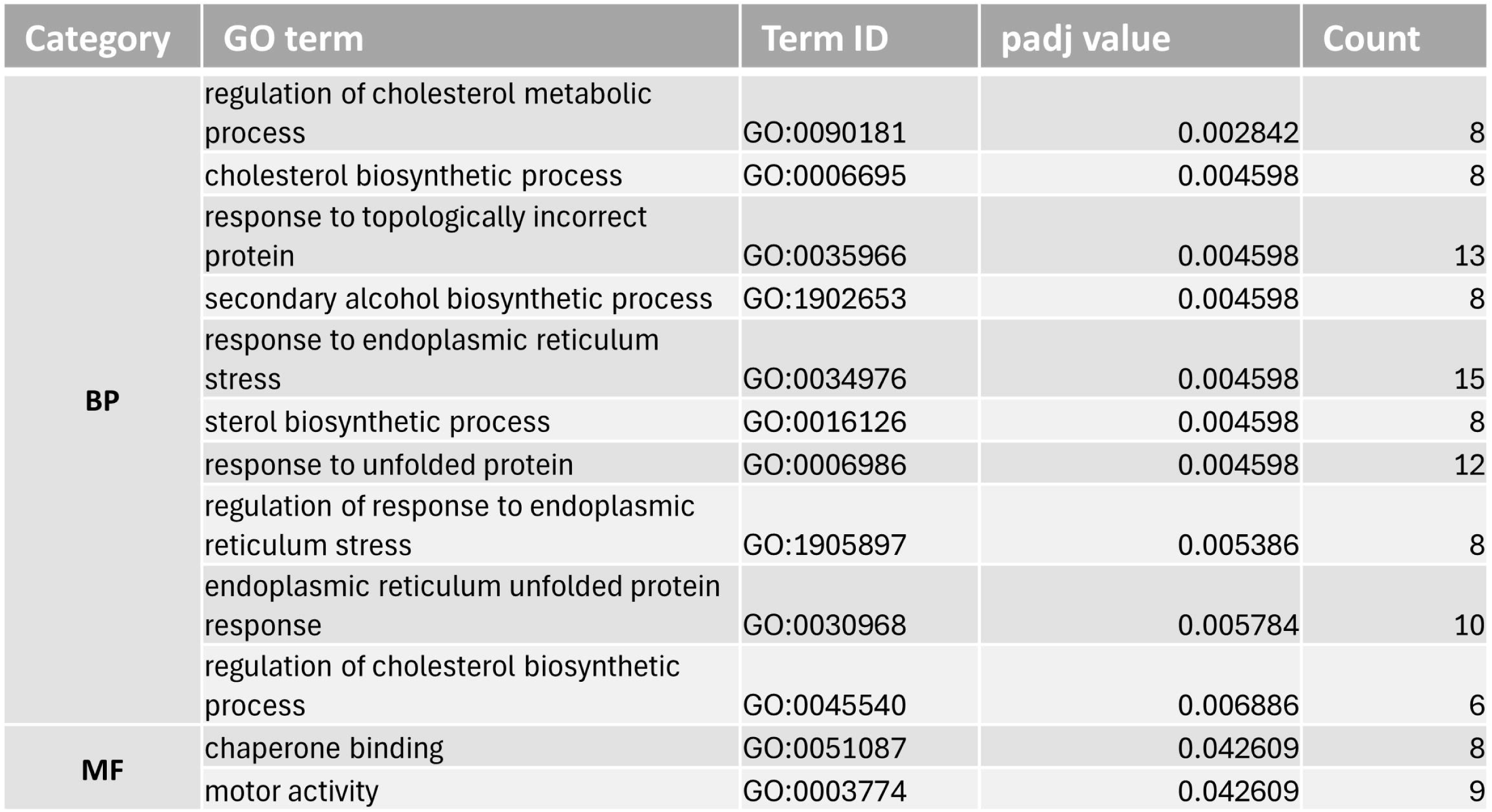
The top 10 significant GO terms for BP and all significant MF GO terms identified through analysis of preferentially enriched CT27 STB DEGs. A comprehensive list of all GO terms can be found in Supplemental File 7.

Finally, both the TSCs and STBs showed differences in the major histocompatibility complex (MHC) molecules. Interestingly, there appears to be an inverse relationship between *HLA-G* and *HLA-C* expression in the CT27 and CT29 TSCs and STBs. *HLA-G* is significantly higher in the CT29 TSCs and STBs, whereas *HLA-C* is significantly higher in the CT27 TSCs and STBs. There was a modest but not significant decrease (padj = 0.07) in *HLA-E* in the CT27 TSCs and a moderate and significant decrease (p < 0.05) in *HLA-E* in the CT27 STBs compared to the CT29 STBs (Figure 5C-H). Collectively, these data demonstrate that there are inherent differences in gene expression between the CT27 and CT29 TSC and STB cell lines.

### TSC cell lines respond differently to IFNB stimulation

We next assessed differences between the CT27 and CT29 TSCs after treatment with IFNB. Gene ontology analysis revealed the top GO terms enriched in the CT29 TSCs were associated with type I interferon signaling and response to virus (Figure 6A, Supplemental File 8). We used hierarchical clustering analysis to compare differences in key genes related to the antiviral, interferon response and found many of these genes were significantly increased in the IFNB-treated CT29 TSCs compared to IFNB-treated CT27 TSCs (Figure 6B). However, these differences were restricted to the TSCs and not observed between the CT27 and CT29 STBs. Most of the pathways enriched in the IFN-treated CT29 STBs compared to the CT27 IFN-treated STBs, were also enriched in the untreated CT29 STBs. These terms were mostly related to the organization of the extracellular matrix, membrane fusion, and epithelial cell proliferation, suggesting differences in differentiation potential between the CT29 and CT27 TSCs.

**Figure 6.**
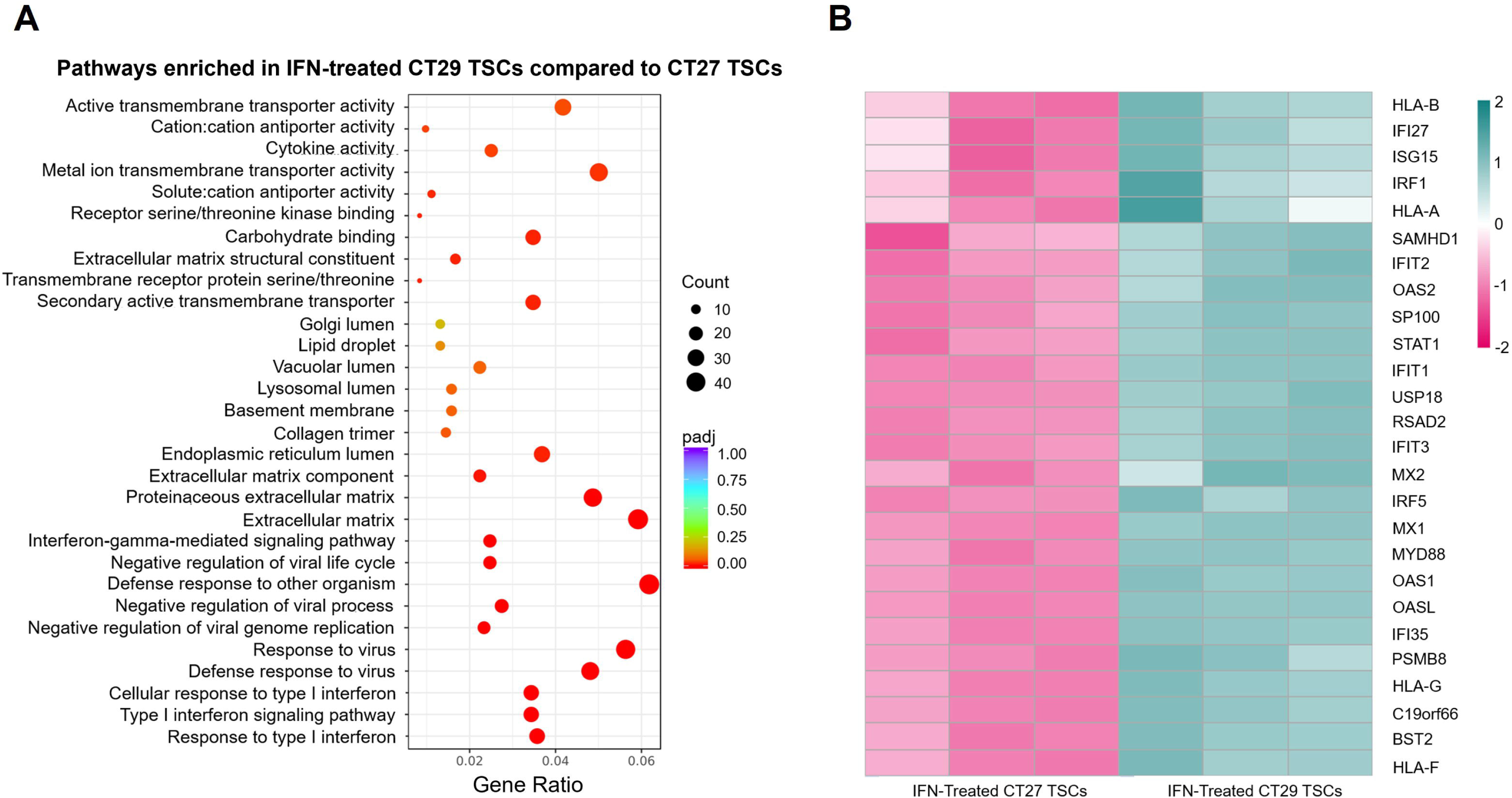
CT29 TSCs have a more robust antiviral response compared to CT27 TSCs. **A)** Gene ontology analysis comparing the IFNB-treated CT29 TSCs to the IFNB-treated CT27 TSCs revealed pathways related to response to virus and type I interferon signaling pathway were significantly enriched in the IFNB-treated CT29 TSCs. Gene ratio refers to the ratio of significant DEGs versus the total number of annotated genes within a specific GO term. **B)** Heat map representation of interferon-stimulated genes and genes related to the interferon signaling pathway. The color spectrum, ranging from teal to pink, indicates high to low normalized levels of expression of each gene.

## Discussion

A balanced maternal immune activation response is paramount to peri-implantation placental development and embryo survival. This study demonstrates an interesting phenomenon where TSCs and STBs do not mount a pro-inflammatory cytokine response in the presence of molecules representative of a microbial infection. Messenger RNA levels of the pro-inflammatory cytokines *IL6* and *IL8* were not increased in either the TSCs or STBs after treatment with LPS. Interestingly, when we performed Western blotting to determine phosphorylation of ERK 15, 30, and 60 minutes after LPS-treatment, we saw a delayed and sustained increase in pERK at 30 and 60 minutes in the TSCs and at 30 minutes only in the STBs. This delayed phosphorylation suggests that the classical TLR4 signaling pathway is present but potentially altered and may become responsive to LPS in alternative circumstances. We believe that our findings support the “double hit hypothesis” which is a hypothesis that proposes that the placenta has a natural, attenuated response to microbial products but maternal viral infections can sensitize the placenta to microbial insults and induce pre-term birth (Cardenas et al., 2011), The studies supporting the double hit hypothesis used human trophoblast cell lines, primary human trophoblast cells, and placentas collected from pregnant mice to demonstrate that placental cells have attenuated pro-inflammatory cytokine responses to LPS. However, pre-treatment of animals and cells with a viral infection sensitized cells and tissues to LPS and lead to a robust cytokine response (Cardenas et al., 2011). Our findings in this manuscript support the hypothesis that the placenta has the ability to recognize LPS through TLR4; however, the response is attenuated without an initial insult the primes that placenta to respond to bacterial products.

We did demonstrate that placental cells are responsive to type I interferons. The role of type I interferons during pregnancy and placental development is complicated and yet to be fully understood. A muted or absent type I interferon response can have detrimental effects on placental development and embryonic and fetal health. Mice lacking the type I interferon receptor (IFNAR) have diminished spiral artery remodeling, albeit the pups were only slightly growth restricted (Murphy et al., 2009). Loss of fetal and placental IFNAR in mice can lead to fetal viremia and increased infection-led fetal mortality compared to wild type mice (Racicot et al., 2017). Additionally an active placental type I interferon response can reduce maternal viral replication in IFNAR mutant dams, suggesting that the placenta offers protection against viremia in the mother (Racicot et al., 2017).

While IFNAR signaling is critical for protection against viral replication in the placenta, fetus, and mother, the detrimental effects of increased IFNAR signaling appears to outweigh the benefit. Women with elevated type I interferon signaling caused by autoimmune disorders and interferonopathies are often at increased risk for maternal morbidity and mortality (Andrade et al., 2015; Gupta & Gupta, 2017; Meuwissen et al., 2016; Ostensen & Clowse, 2013). In the context of an active infection, Zika and other viruses induce a potent type I interferon response that disrupts placental development and induces embryonic and fetal demise (Miner et al., 2016; Yockey & Iwasaki, 2018; Yockey et al., 2018). IFNAR signaling in response to Zika causes increased apoptosis in the labyrinth layer of the murine placenta and aberrations at the maternal-fetal blood barrier leading to mixing of maternal and fetal blood within the labyrinth (Yockey et al., 2018). The labyrinth layer of the murine placenta is analogous to the STB layer of the human placenta. Interestingly, treatment of chorionic villous explants with IFNB led to the formation of syncytial knots and significant decrease in human chorionic gondatotropin subunit beta (CGB) mRNA levels (Yockey et al., 2018). Collectively, these previous studies suggest that most of the disruption of placental development and function occur at the STB layer.

We demonstrate that IFNB induces a robust increase in *IFITM1*, *IFITM2*, and *IFITM3* in both TSCs and STBs. This phenomenon has also been reported in immortalized choriocarcinoma cells and term villous cytotrophoblast cells (Buchrieser et al., 2019). Additionally, high levels of the IFITM proteins inhibit STB formation and trophoblast cell invasion (Buchrieser et al., 2019; Degrelle et al., 2023; Zani et al., 2019), which can induce fetal demise. As the TSCs and STBs we used are molecularly representative of the peri-implantation stage placenta, we propose IFNB induction of IFITM proteins may activate a “safety switch” mechanism to prevent the finalization of implantation and adequate trophoblast invasion in a hostile uterine environment (West et al., 2019). This molecular safety switch may serve to spare the potentially compromised mother from expending energy on the growth of an embryo and eventual fetus.

To comprehensively analyze transcriptional differences in IFNB stimulation between TSCs and STBs, we used a 3D spheroid model of immune stimulation in the TSCs and STBs. We opted to use a spheroid model for transcriptomics as there has been speculation that the coating substratum used to facilitate cell attachment on tissue culture-treated plates might be influencing the gene expression of TSCs (Logsdon et al., 2024; West et al., 2019). By allowing the TSCs and STBs to grow and organize in suspension culture, we removed the influence of the coating substrate. Transcriptional data comparing the IFNB-treated TSC spheroids and IFNB-treated STB spheroids revealed key differences in how the cell types mount an antiviral response. While both TSC and STB spheroids respond to IFNB, the untreated STB spheroids seem primed to produce a more robust and coordinated antiviral response, potentially due to their role as the primary maternal-fetal barrier that protects the fetus from maternal-fetal transmission of pathogens. Our results suggest that TSCs and STBs can induce distinct antiviral responses, with STBs upregulating pathways that reflect antigen sensing and communication with the uterus. As the STB is the layer of the placenta that comes into direct contact with the endometrium during invasion, these data are consistent with the idea that the placenta can provide antiviral factors to assist the maternal immune response in protection against maternal viremia (Racicot et al., 2017).

We also provide evidence that there are inherent differences between the commonly used CT27 and CT29 TSC lines. Of relevance to the placental immune response, there were significant differences in the expression of the human leukocyte antigen (HLA) molecules, *HLA-C, HLA-E,* and *HLA-G*, between the CT27 and CT29 TSCs and STBs. While *HLA-C* was significantly reduced in both CT27 TSCs and STBs compared to the CT29s, *HLA-E* and *HLA-G* were significantly increased. Previous studies have reported increased levels of HLA-G in EVTs and in amniotic fluid from pregnancies carrying a male fetus (Emmer et al., 2003; Papuchova, Meissner, Li, Strominger, & Tilburgs, 2019), suggesting that there is a sexual dimorphism to the expression of HLA molecules in extraembryonic tissues. We also observed sex-specific differences in the MT gene family. Sexually dimorphic expression of MTs has been described in other tissues, with MT expression increased in female livers and kidneys (Kim, Kim, Park, Ryu, & Yu, 2009; Zhang et al., 2012). Additionally, in a pregnant mouse model, Cadmium (Cd) exposure led to significantly elevated levels of *Mt1* and *Mt2* in female placentas compared to male and female fetal livers had five times the accumulation of Cd compared to male fetal livers, indicating increased transfer of metals across the placenta in females (Jackson, Baars, & Belcher, 2022).

There were also differential responses to IFNB treatment between the CT27 and CT29 TSCs. The CT29 TSCs seemingly exhibit a stronger interferon response, with several ISGs and genes related to the IFN-signaling pathway significantly higher in the IFN-treated CT29 TSCs compared to the CT27 TSCs. Our results are consistent with previous findings that male placentas have a heightened ISG response after viral exposure (Bordt et al., 2021); however, these differences were not observed in the STBs.

We recognize that one shortcoming of this study is the small sample size. By only comparing the CT27 and CT29 TSCs, it is difficult to determine if the observed differences were caused by inherent genetic variation or sex. While we did observe several sex-specific phenomena that have been described in other tissues in our cell lines, further research with a larger sample size is necessary to make more substantive conclusions. Our main purpose in comparing the cell lines was to identify differences that potentially make one cell line better than another to address different hypotheses related to the placental innate immune response. Our data also reinforce the inclusion of sex as a biological variable for in vitro work. Future work should address the limited sample size in our study by including sex in the analyses of clinical samples or in vivo animal models.

Our results reinforce a growing body of work demonstrating the complexity of the peri-implantation placental immune response. Implantation failure is often attributed to an unknown etiology; however, the role of the maternal immune milieu can no longer be overlooked as a potential culprit. By responding to IFNB but not LPS, the TSCs and STBs underscore the importance in the host response to microbial infections during pregnancy. Better understanding how type I interferons disrupt placental development, especially during the peri-implantation period, creates new insight into how the uterine immune response influences embryo health. This molecular investigation of the peri-implantation period using representative stem cell models implicates the primitive placenta as a key mediator of maternal immune balance and opens new avenues to unlock the black box of early pregnancy loss.

## Declaration of interest

RCW is an Associate Editor for *Reproduction & Fertility* and was not involved in the review or editorial process for this paper.

## Funding

This research was supported by the Ky Cha Award in Stem Cell Technology awarded by the American Society for Reproductive Medicine and the Auburn University Animal Health and Disease Research Fund.

## Author contribution statement

CRC performed experiments, contributed to study design, contributed to data analysis, and critically reviewed the manuscript. JB, MB, PD, and CP conducted the experiments. RCW conceived the study, performed data analysis, secured research funding for the study, and wrote the manuscript.

## Acknowledgments

We would like to acknowledge Victoria Caravaggio (Auburn University) for her assistance on the project. We would also like to thank Dr. Mihnea Mangalea (CDC) for his critical review of the manuscript.

---

**Supplemental Figure 1.**
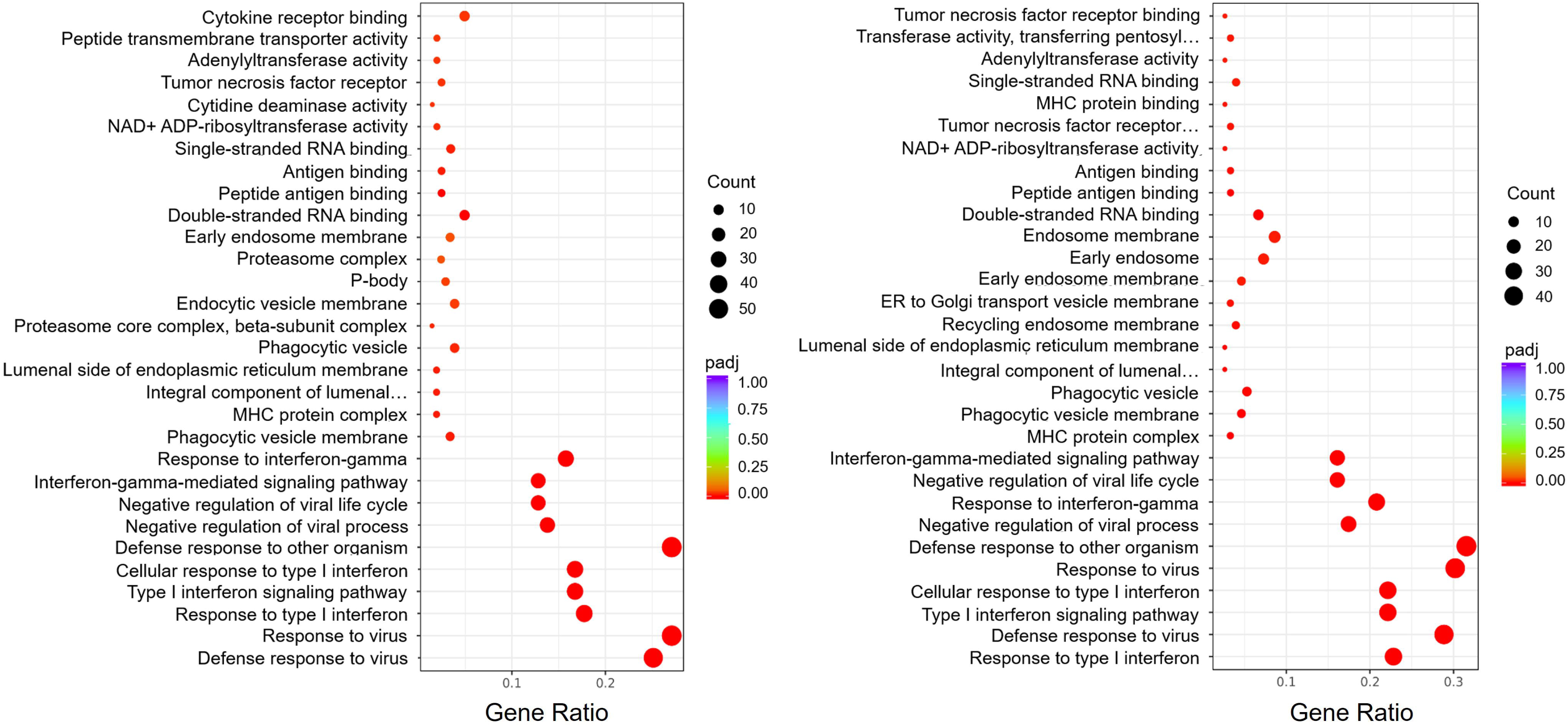
Gene ontology and pathway analysis of IFNB-treated TSCs vs. TSCs and IFNB-treated STBs vs. STBs. The pathways enriched in IFNB-treated TSCs and STBs are similar, with pathways related to defense response to virus and response to type I interferons being the most significantly enriched in both cell types compared to untreated control cells.

**Supplemental Figure 2.**
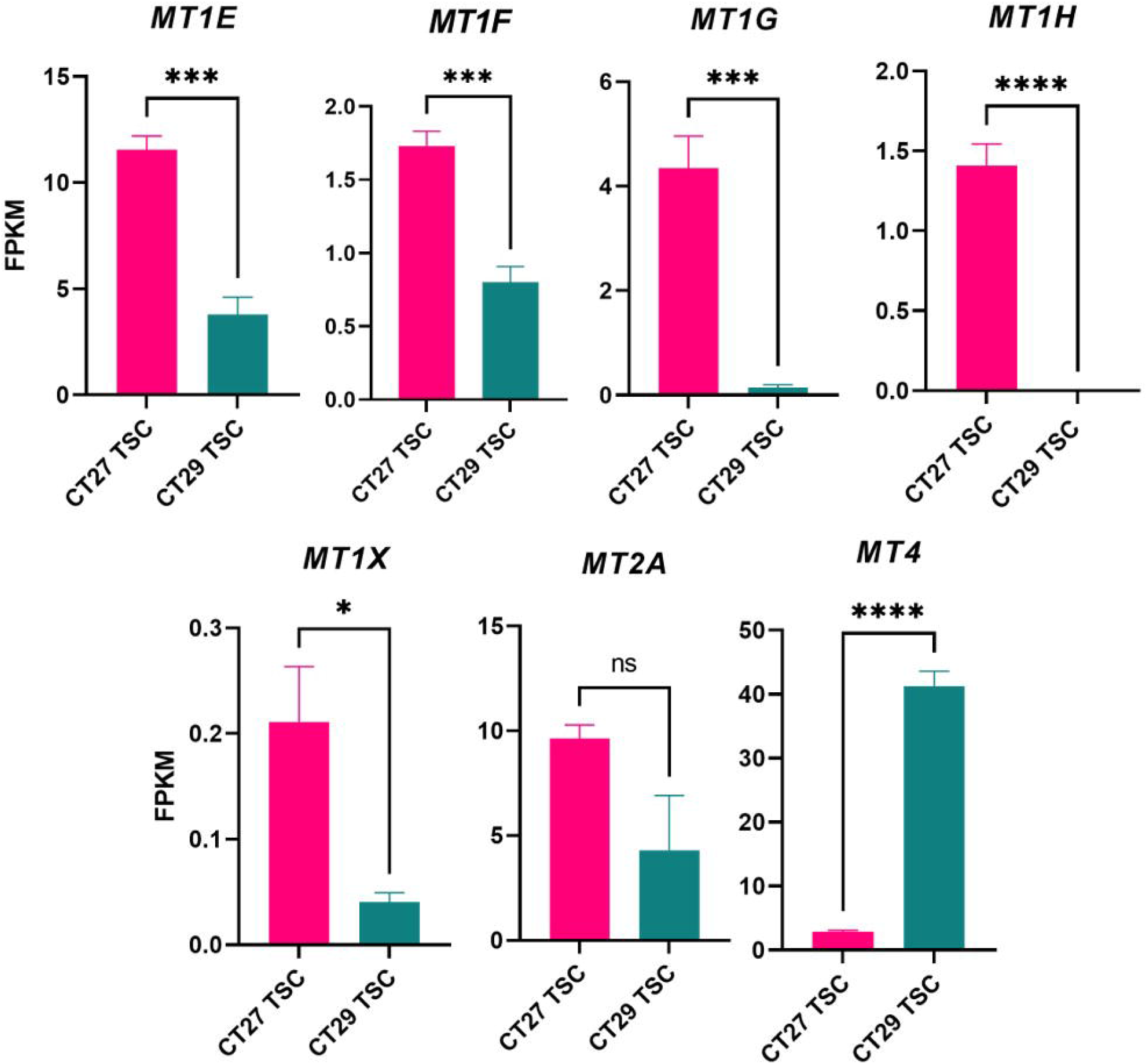
Genes belonging to the metallothionein pathway are significantly higher in the CT27 TSCs compared to the CT29 TSCs. Relative expression levels of metallothionein genes. The values are FPKM values and error bars represent the SEM calculated from the RNA-seq analysis. * indicates padj < 0.05, *** indicates padj < 0.005, **** indicates padj < 0.0001.

## Notes

### Competing Interest Statement

The authors have declared no competing interest.

### Summary of Updates

This version of the manuscript has been revised to update the RNAsequencing analysis.

